# Spontaneous thought orientation tracked by fMRI networks and EEG alpha power dynamics

**DOI:** 10.1101/2025.10.31.684884

**Authors:** Tomas Hampejs, David Tomecek, Stanislav Jiricek, Vlastimil Koudelka, Lucia Jajcay, Dante Mantini, Jaroslav Hlinka

## Abstract

Understanding how spontaneous, rather than experimentally induced, thoughts relate to brain activity remains a major challenge. We combined simultaneous fMRI and EEG recordings with Descriptive Experience Sampling (DES) to link momentary, naturally occurring experiences to their neural signatures during rest. Using machine-learning classification of 240 time-locked samples from eight participants—each completing nine 25-minute resting-state sessions—we reliably distinguished internally from externally oriented experiences (fMRI accuracy = 65.4%, EEG = 62.5%). Externally oriented states showed greater fMRI activity in salience, auditory, and visuospatial networks and lower occipital alpha power in EEG, whereas internally oriented states exhibited the opposite pattern, extending prior DMN-focused accounts of internally directed states. Across modalities, integrated resting-state alpha power correlated negatively with BOLD fluctuations in parietal and occipital regions. These multimodal findings reveal distinct neural signatures of spontaneous experience and demonstrate that coordinated large-scale network dynamics and alpha-band oscillations track the natural alternation between inward and outward focus in the human mind.

## 1 Introduction

Human behaviour research has continuously evolved in its core questions and methodological approaches. Early psychology in the nineteenth century was marked by its reliance on introspection—self-observation—as a primary scientific tool [1, 2]. For William James [3], introspection provided privileged access to the “stream of consciousness,” the continuous flow of mental states that formed psychology’s central object of study. In contrast, Wilhelm Wundt advocated a more systematic, experimental introspection aimed at measuring mental processes with precision [4]. Over time, it was Wundt’s controlled experimental approach—not James’s phenomenological one—that shaped mainstream psychology. Behaviourism subsequently displaced explicit introspection, replacing it with observable responses or standardized questionnaires, and even the rise of cognitive psychology largely retained this emphasis on task-based experimentation. Throughout these paradigm shifts, the link between mental states and their physiological underpinnings remained a persistent theme. Electroencephalography (EEG), since its invention by Hans Berger in 1924 [5], became a cornerstone for investigating neural correlates of both externally triggered and spontaneous brain activity. Its oscillatory rhythms, particularly the waxing and waning parieto-occipital alpha waves, were associated with relaxed wakefulness and internally directed states.

The introduction of functional magnetic resonance imaging (fMRI) in the early 1990s [6] revolutionized human brain mapping, initially focusing on localizing functions through controlled experimental designs. A major conceptual leap followed with the recognition that even in the absence of explicit tasks, fMRI reveals coherent spontaneous fluctuations in brain activity—the so-called resting state. These fluctuations display reproducible patterns of functional connectivity [7, 8], correlate with concurrent EEG activity [9–13], and vary meaningfully across behavioural and clinical states. Among these large-scale systems, the *default mode network* (DMN) [14, 15] is most consistently linked to internally focused cognition such as self-referential thought and memory retrieval. The DMN alternates in activation with the *central executive network* (CEN) [16], while the *salience network* mediates switching between internally and externally oriented attention [17]. Externally oriented processing relies on the *visuospatial* or *dorsal attention network* (DAN) [18–20]. Together, these networks form a dynamic system through which the brain flexibly transitions between perceptual engagement and internally generated mentation [8].

This technological and conceptual progress has revived long-standing philosophical questions about linking neural activity to subjective experience. Despite occasional metaphysical [21] or ethical concerns [22, 23], the key challenge remains empirical: integrating neuroimaging’s complementary spatial and temporal strengths with ecologically valid measures of lived experience. While the DMN–CEN/DAN framework contrasts internal and external processing, it relies mainly on inferences from task-based activations and thus remains partly speculative. A few pioneering studies attempted to probe momentary experience during resting-state fMRI [24–27]. Although informative, they faced limitations in temporal precision and veracity of experiential sampling. EEG, with its superior temporal resolution, offers closer access to rapidly changing mental events, yet no previous EEG study has explicitly examined internally versus externally oriented streams of thought under resting conditions. Task-based EEG research consistently links higher alpha power to internally directed attention (e.g., imagery, mental arithmetic) and lower alpha power to externally focused states [28–30]. Mind-wandering studies confirm this pattern, showing increased alpha power during internally focused or off-task moments [31–34]. Accordingly, fluctuations in alpha power during rest are expected to mark shifts between inward and outward attentional orientation. Alpha oscillations are particularly valuable because their spectral dynamics covary with fMRI BOLD fluctuations across widespread cortical areas [9, 11–13, 35–39], thereby bridging the modalities. Recent methodological advances [13] allow extraction of robust spatio-spectral EEG patterns, including the parieto-occipital alpha configuration reliably associated with BOLD signal changes. This multimodal complementarity provides a unique window into the spatiotemporal dynamics underlying spontaneous experience.

Yet, despite substantial progress in acquiring brain activity data, conventional multimodal resting-state neuroimaging still falls short in incorporating a “phenomenological modality,” that is, in systematically capturing comparatively rich data about participants’ mentation. Traditional post-hoc questionnaires [40, 41] offer only coarse, retrospective summaries and are prone to introspection bias [42–44]. To overcome these limitations, *experience sampling* (ES) techniques prompt participants at random moments to report their immediate experience [45, 46]. ES increases temporal precision and ecological validity but varies widely in implementation — from closed-choice categorizations to open-ended and “open-beginniged” phenomenological reports [47–51]. The most rigorous of these approaches, Descriptive Experience Sampling (DES) [51], minimizes framing bias by using brief auditory cues (“beeps”) and elicitation interviews to reconstruct “pristine” moments of awareness. Several fMRI studies have combined ES with neuroimaging to contrast internal and external awareness [24–27, 52]. Typically, internally oriented states engage the DMN, whereas externally oriented ones activate attentional networks. However, methodological differences limit comparability: some studies required participants to classify fixed temporal windows (e.g., 10 s [26], several seconds [25]), while others allowed variable durations up to 30 s [27]. Frequent probing can itself create an implicit monitoring task [24], compromising the resting-state condition. Moreover, except for Fernyhough et al. [24], most studies imposed discrete a priori categories (thought vs. perception), neglecting the nuanced continuum of experience.

Fernyhough et al. [24] addressed these issues by combining fMRI with DES, enabling post-hoc consensus coding of experiential sample descriptions based on elicitation interviews. Despite its small sample (five participants), the study set a benchmark for integrating rigorous phenomenological methods with neuroimaging, demonstrating the feasibility of linking spontaneous inner experience with brain activity.

Building on these advances, the present study aims to extend our understanding of the brain–mind relationship by identifying neural correlates of internally and externally oriented spontaneous awareness using state-of-the-art multimodal tools. We combine resting-state fMRI with Descriptive Experience Sampling [51] to capture the richness of naturally occurring mentation, replicate and expand the study of Fernyhough et al. [24] on a larger sample with modern acquisition, and complement it with simultaneous EEG recordings for superior temporal resolution and cross-validation. Beyond traditional statistical analyses, we employ a machine learning framework to classify brain states based on experiential states, thereby enhancing interpretability and facilitating the detection of complex spatiotemporal patterns. This first report focuses on the fundamental contrast between internally and externally oriented experience, providing a foundation for future analyses that exploit the full methodological potential of the current multimodal dataset. By pairing DES with simultaneous EEG–fMRI, we link first-person reports to third-person neural dynamics within the same moments of rest. This integration leverages the complementary strengths of BOLD-based large-scale network mapping and alpha-band electrophysiology to capture how the brain alternates between inward and outward modes of awareness. In doing so, we move beyond task-derived inference about internal versus external cognition and toward direct, time-resolved correspondence between experience and brain activity. The present study thus positions spontaneous thought not as measurement noise, but as an organized, multimodal signature of ongoing cognition, setting the stage for mechanistic accounts of how large-scale networks and oscillatory rhythms shape the natural ebb and flow of human experience.

## 2 Results

### 2.1 Descriptive Experience Sampling and Data Overview

Across eight participants, we collected data in nine resting-state EEG–fMRI sessions per subject (25 min each). While one session contained four sampling events, we gathered in total 288 DES samples (descriptions of pristine momentary experience states). From these, 240 reached full coder consensus over target category (163 “internally oriented”, 77 “externally oriented”) and were retained for analysis (Tab. 1); 48 samples were excluded due to disagreement (41) or motion artefacts (7) in corresponding brain activity data.

**Table 1:** Number of internal/external (I/E) samples per subject (names are pseudonyms).

| Subject | Internal | External |
| --- | --- | --- |
| Jane | 22 | 9 |
| Case | 21 | 6 |
| Eva | 27 | 6 |
| George | 21 | 9 |
| Anna | 14 | 13 |
| Nika | 18 | 15 |
| Valerie | 29 | 3 |
| Tamara | 11 | 16 |

### 2.2 Multimodal Classification of Internal and External Experience

We quantified neural separability between internally and externally oriented experience using machine-learning classification on both fMRI and EEG data (Fig. 1). Separate linear SVMs were trained on time-locked features from each modality, with accuracy used as an empirical measure of distinction. This approach integrates signals across networks rather than testing them individually, providing a direct estimate of how reliably spontaneous experiential states can be decoded from brain activity.

**Fig. 1:**
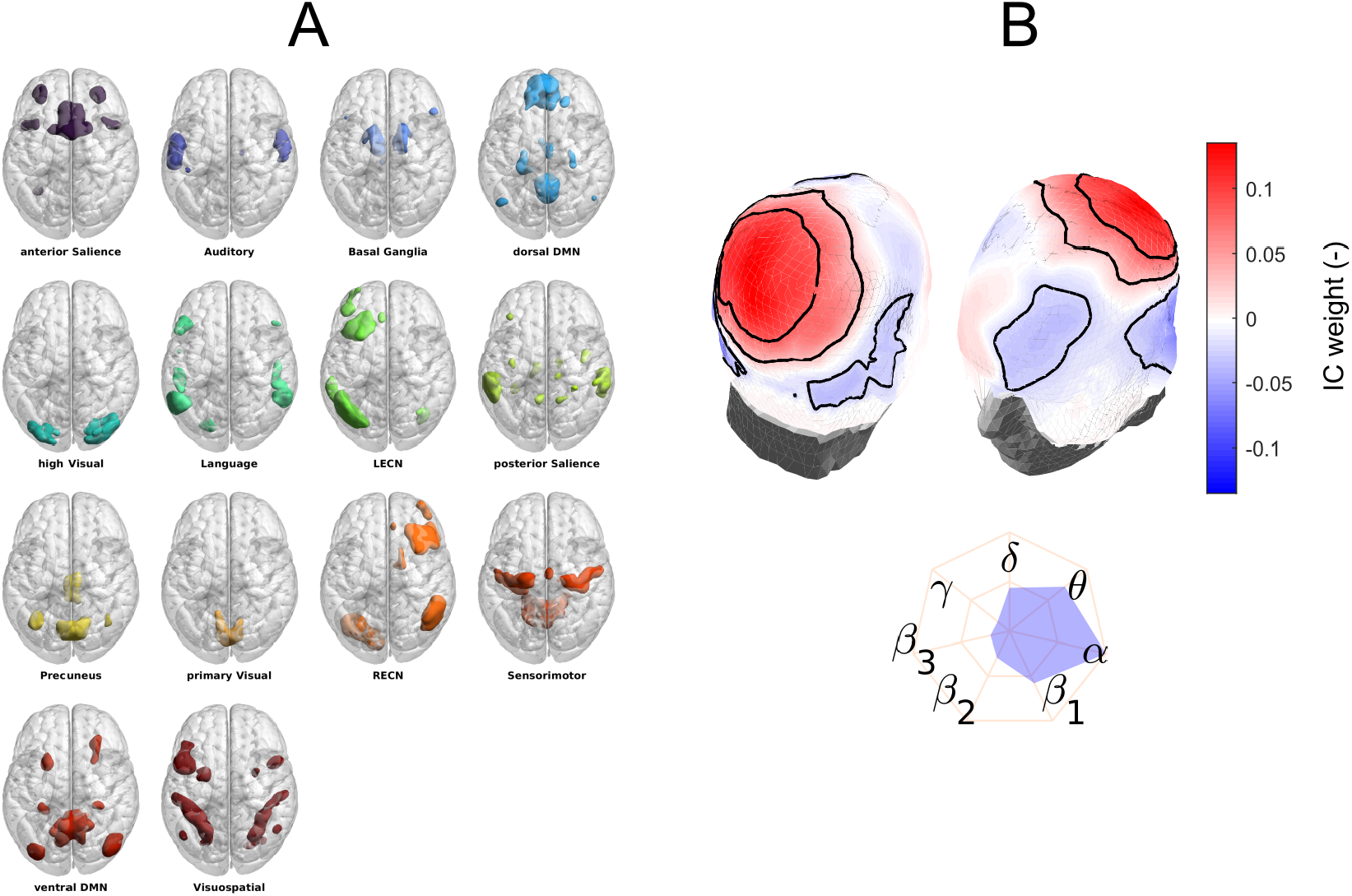
Feature extraction from simultaneous EEG–fMRI. **(A)** Fourteen resting-state network masks from [53]. **(B)** EEG *α* spatio-spectral pattern from [13].

A combined EEG–fMRI classifier reached 65.0 % accuracy—comparable to the fMRI-based model—suggesting that most discriminative information resides in the BOLD features. Consequently, the subsequent analyses focus on modality-specific classifiers.

#### 2.2.1 BOLD-based Classification

Using time series from 14 functional networks [53], the BOLD-based classifier achieved statistically significant discrimination between I/E states (TFCE, *p <* 0.05), peaking at 65.4 % accuracy at multiple positions, reflecting the temporal smoothness of the input BOLD signals and the averaging of identical BOLD samples across multiple window parameter combinations. For instance, a maximum at a window size of 0.1 s and a window position of −4.1 s before the beep (Fig. 2B), with a balanced accuracy of 64.3%, correctly identified 110 of 163 internal and 47 of 77 external samples, as illustrated by the confusion matrix in Fig. 2B.Identical accuracies were also observed at a window size of 0.4 s (position −2 s), 1.8 s (position−3.2 s), and 1.5 s (position −3.3 s).

**Fig. 2:**
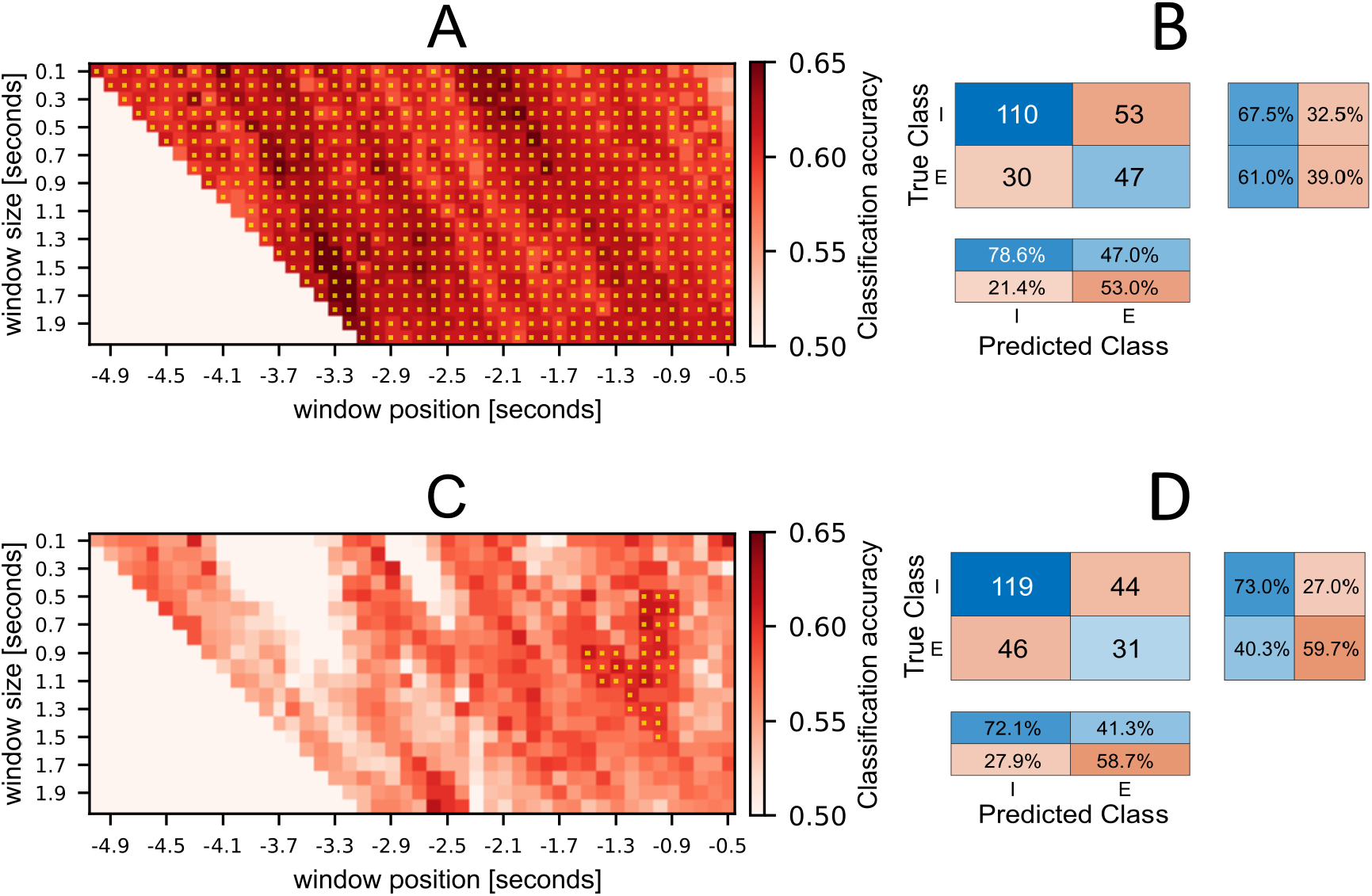
Classification performance for BOLD (**A–B**) and EEG (**C–D**) data. Yellow regions in **A, C** indicate significant TFCE clusters (*p <* 0.05).

#### 2.2.2 EEG-based Classification

EEG analyses focused on the *α* spatio-spectral pattern—an index of attentional gating [13]. Of 30 independent components, two parietal *α* sources were merged to form a single temporal feature (Fig. 1B). Classification based on its instantaneous amplitude reached a maximum accuracy of 62.5% (balanced = 56.6%) for a window size of 0.7 s and a window position of −1.1 s, correctly predicting 119 internal samples out of 163 and 31 external samples out of 77 (see the confusion matrix in Fig. 2D). The cluster (TFCE, *p <* 0.05) with the most successful classification occurred within windows from −1.5 to −0.9 s before the beep and with durations of 0.5–1.5 s (Fig. 2C). The temporal window with substantial EEG discrimination was generally more confined than in fMRI, consistent with the expected finer sensitivity to pre-conscious neural transitions.

An additional analysis using non-averaged 100ms EEG samples was conducted to test whether temporal averaging obscured fast dynamics. The accuracy did not substantially differ from the best EEG classifier (60.7 %, p = 0.02), indicating that averaging did not inflate performance.

### 2.3 Exploratory Time-course Analysis

To assess broader temporal dynamics, we examined 15s windows around the sampling event (Fig. 3). Five BOLD networks—anterior and posterior Salience, Auditory, high Visual, and Visuospatial—showed the strongest internal/external differences (Wilcoxon, *α* = 0.01). These patterns largely aligned with the 5s pre-event classification window, implying consistent pre-sampling network reconfiguration.

**Fig. 3:**
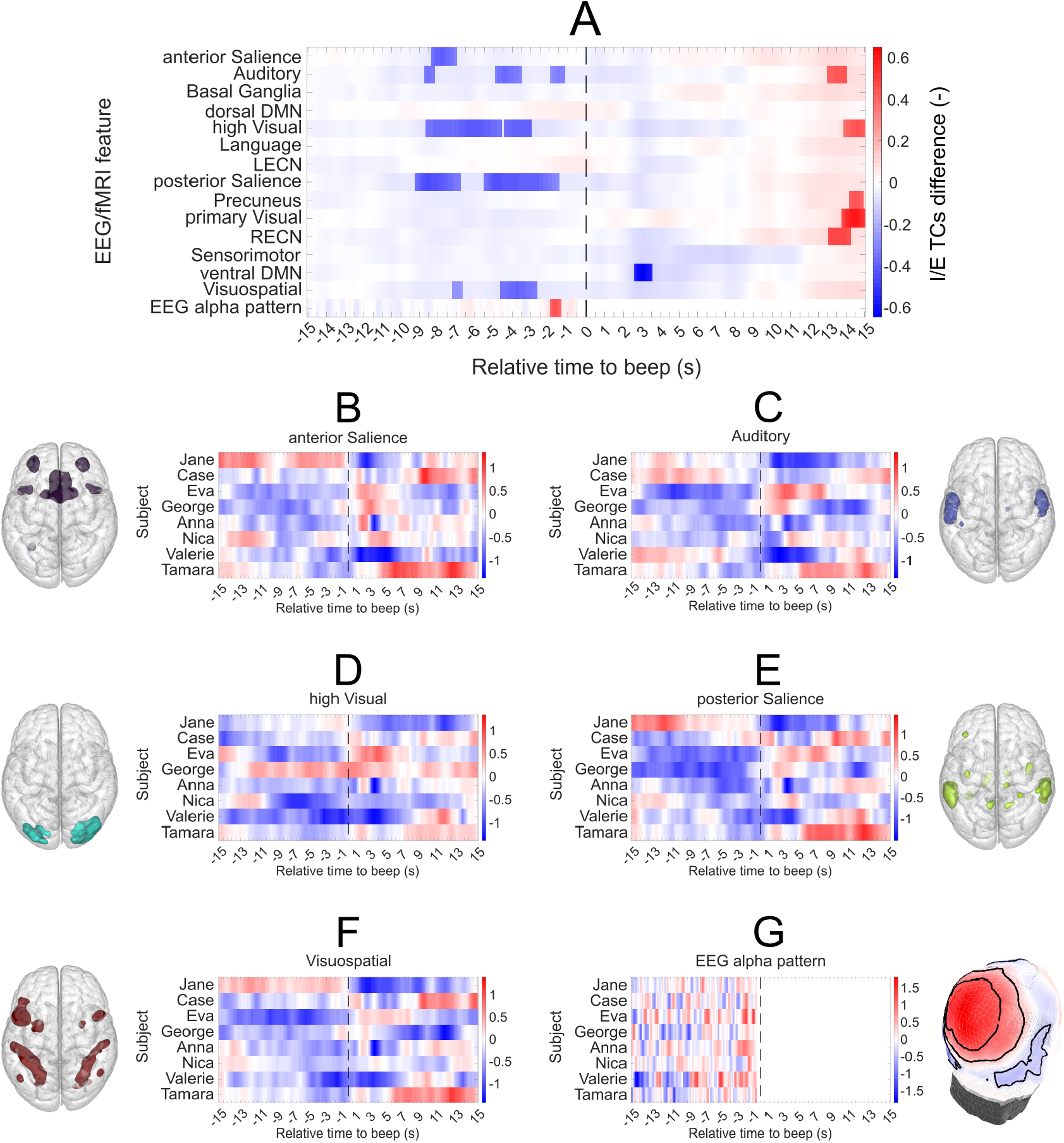
Time-resolved I/E differences in BOLD and EEG features. Opaque segments indicate significant intervals (*α* = 0.01, Wilcoxon).

Single-subject traces revealed shared tendencies but also notable variability: for example, Jane displayed inverted differences in three networks, and George in the high-visual network. Similarly, EEG *α* differences were generally consistent except for one participant (Nika), whose polarity reversed between –2 and –1.5s.

Node-wise exploration of these five networks (Fig. 4) identified key contributors within the inferior parietal lobule, visual cortex, and insula—regions known for multimodal integration and attentional control. This supports the view that sensory and salience systems dominate the neural signature of externally oriented experience. On the other hand, in contrast to prior work, no reliable differences were observed within DMN nodes, suggesting that while the DMN is in stark contrast with common attention-demanding tasks, it is not strongly involved in the contrast between internally and externally oriented unconstrained mentation. Of course, this result may be modulated by the contribution of specific types of mentation present in the subjects during the experiential sampling [54].

**Fig. 4:**
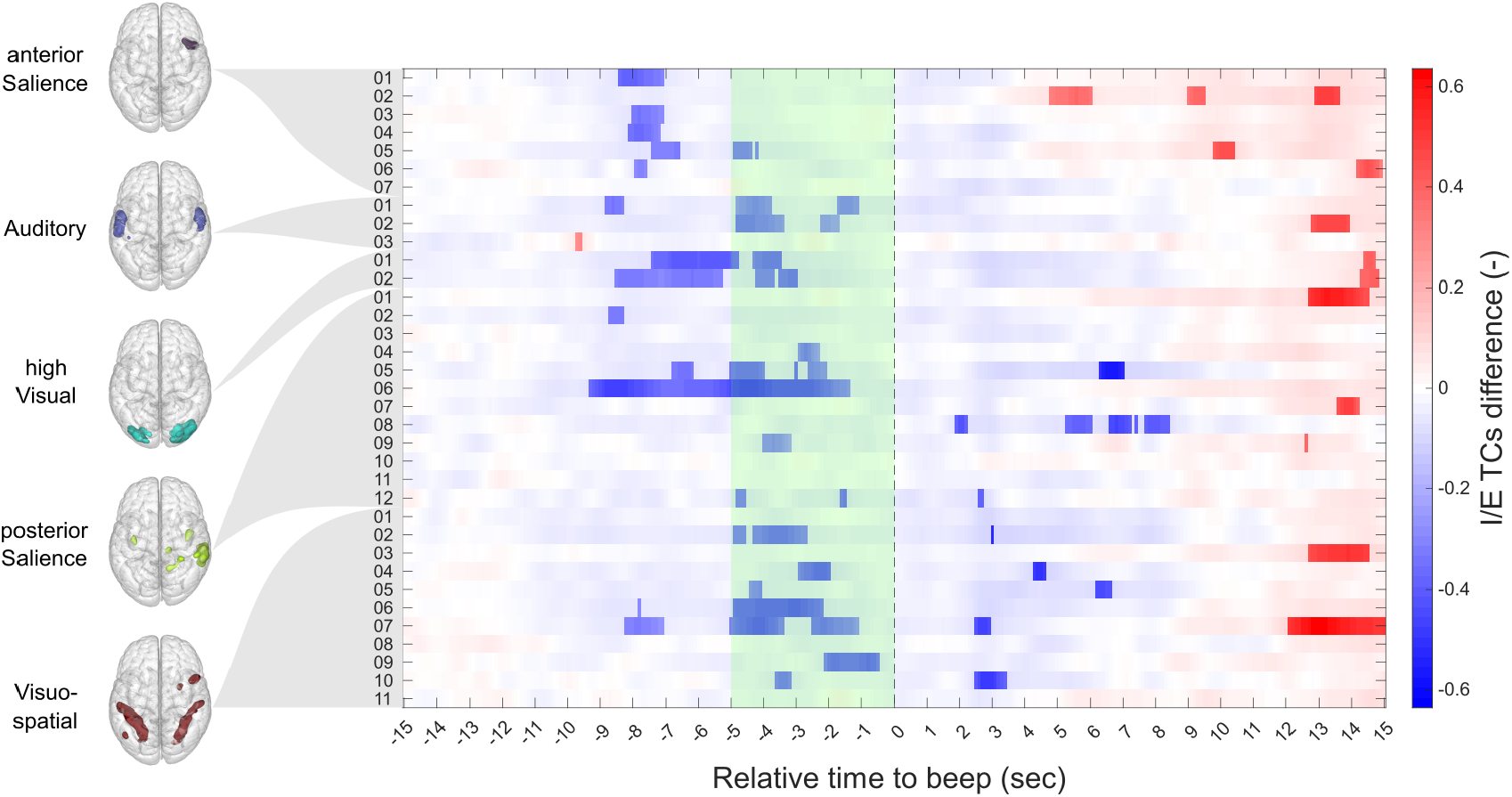
Node-wise BOLD exploration for five significant networks. Opaque segments mark significant intervals (*α* = 0.01, Wilcoxon).

### 2.4 Coupling between EEG *α* and BOLD Fluctuations

To validate the electrophysiological marker, we evaluated voxelwise correlations between the EEG *α* spatiospectral amplitude and the BOLD signal during undisturbed rest (first 500 TRs of each run). The resulting group-level GLM map (Fig. 5A) revealed a bilateral negative association in parietal and occipital cortices, replicating earlier findings [13]. A spatial correlation with 14 RSN masks (Fig. 5B) confirmed the strongest spatial overlap with the high-visual network, reinforcing that low *α* power coincides with enhanced sensory processing.

**Fig. 5:**
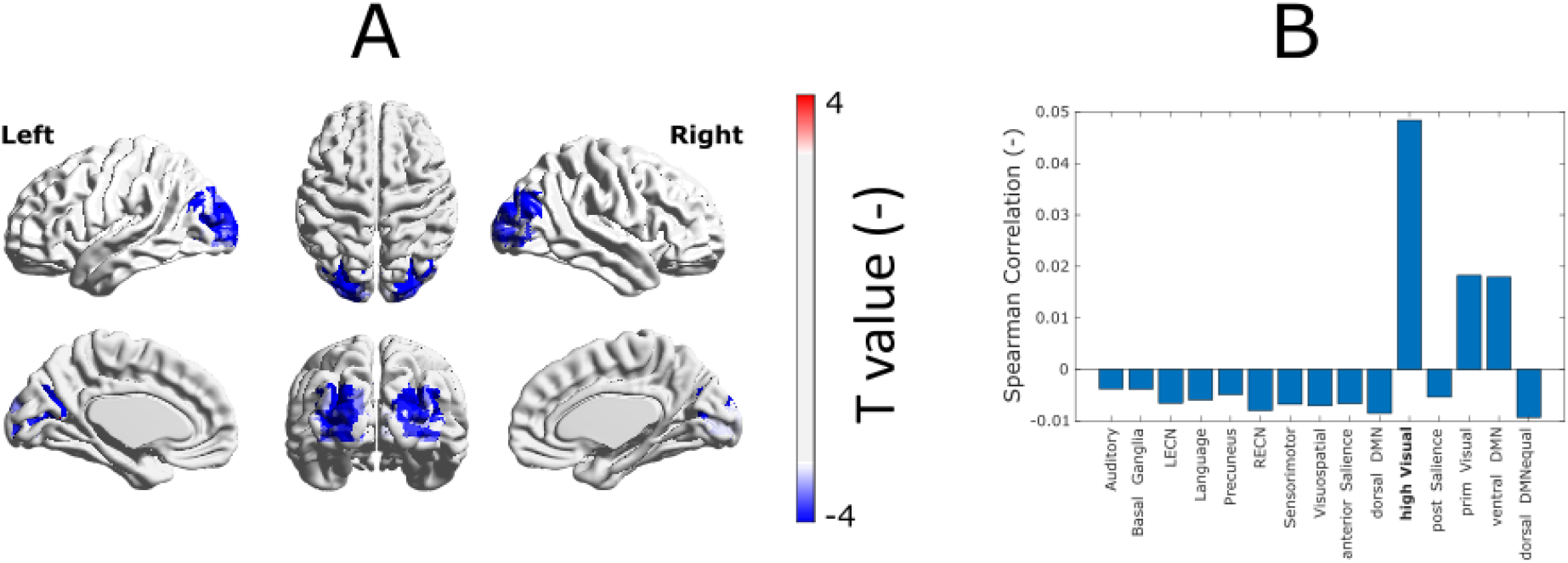
EEG–fMRI integration. **(A)** GLM map of *α* amplitude vs. voxel-wise BOLD (*p*_*cluster*_ *<* 0.05). **(B)** Correlation of the resulting mask with 14 RSNs.

## 3 Discussion

We present an interdisciplinary investigation of the mind–body relationship that combines multi-modal neuroimaging, momentary experience sampling, and machine learning. Our goal was to identify neural differences between internally and externally oriented *spontaneous* experiences during restingstate EEG–fMRI. A quasi-random “beep” paradigm yielded 288 DES samples; 240 (163 internal, 73 external) were reliably classified and used to train models linking experiential states to brain activity.

The reliable discrimination of internal versus external experience from simultaneous EEG and fMRI signals demonstrates a robust link between spontaneous mentation and large-scale neural dynamics [55]. Beyond methodological validation, this supports the view that spontaneous thought is not stochastic ‘noise’ but an organized cognitive mode embedded within intrinsic network architecture [56]. Building on Fernyhough et al. [24], we anticipated DMN engagement for experiential states decoupled from perception, consistent with mind-wandering and its “freedom of immediacy” [57]. Although we did not replicate a DMN effect, we observed the reverse contrast (*external > internal*), suggesting that perceptual and attentional processes within sensory and salience systems, besides the canonical DMN modulation, also distinguish inward and outward modes of experience.

Using machine learning, we classified brain states from time series within 14 networks defined by Shirer et al. [53]. Departing from prior work that used either a single time point [24] or a wide blocks [27, 58] of brain activity data correlated to subjective reports, we systematically scanned window sizes and positions and found surprising uncertainty in anchoring the experiential moment to “brain moment”.

Although DES anchors experience to “the moment just before the beep,” some samples, especially due to its reductive coding, likely reflect broader states unfolding earlier. Exploratory analyses over a wider 15 s pre-beep interval identified consistent group-level differences, with the strongest internal/external contrasts in five networks—anterior salience, auditory, high visual, posterior salience, and visuospatial [53]—all showing greater activity during external states. Notable exceptions (e.g., George: high visual; Jane: posterior salience, visuospatial) highlight meaningful individual variability and caution against over-generalization from single-subject patterns. While DES anchors experience to “the moment just before the beep,” it is likely that samples tend to reflect broader lasting states.

### 3.1 Network-level interpretation

The visuospatial/dorsal attention network (DAN) [18] was prominently engaged during externally oriented experience, consistent with its role in goal-directed behaviour and visuospatial processing [54]. Even without marked DMN deactivation, the well-established anticorrelation between DAN and DMN [16, 54] supports the interpretation that externally oriented states are dominated by attention-to-perception mechanisms rather than by the suppression of internal mentation.

We also observed co-activation of DAN and auditory regions during external states, consistent with *multisensory* recruitment when attention is directed outward [59, 60]. Although only 16 of 77 external samples reported explicit auditory content (and 25 of 163 internal samples included inner hearing), auditory cortex activity can be driven by imagery or inner speech [61, 62]. The observed secondary visual cortex (V2) [63] enhancement during external focus suggests heightened selectivity for relevant inputs and suppression of distractors [64, 65], which supports object recognition and visual memory [66] and aligns with bottom-up recruitment via the DAN [67].

Posterior salience network (pSN) differences, particularly within inferior parietal lobule (IPL), further differentiated externally from internally oriented samples. The pSN supports multimodal integration and sensorimotor linkage [68], consistent with the frequent visual (43 of 77) and bodily (54 of 77) components in external samples. The IPL’s role in visuospatial attention and object manipulation [20, 69] reinforces its contribution; node-level patterns align with Vanhaudenhuyse et al. [27], who reported increased right-IPL activity during external awareness.

The anterior salience network (aSN) contributed to internal–external contrasts, but its peak activity occurred outside the 5 s classification window. As a gatekeeper between salient external stimuli and internal cognition [70–72], the aSN may operate on slower timescales, coordinating transitions rather than moment-to-moment content. Together, these findings indicate a coordinated interplay among DAN, pSN/aSN, auditory, and high-visual networks in regulating spontaneous attention—complementing prior evidence that internally directed states engage DMN [24].

### 3.2 Electrophysiological dynamics of spontaneous attention

Concurrently recorded EEG showed that parieto-occipital *α* power—a hallmark of attentional gating—distinguished internal from external experience with peak classification within a brief (*<*2 s) pre-beep window (Fig. 3A). This is consistent with studies linking *α* to top-down sensory modulation [28, 73–75] and extends them to *spontaneous*, non-instructed states: low *α* implies heightened cortical excitability and outward focus, whereas high *α* implies sensory suppression and inward focus [76]. The temporal compactness of the EEG effect relative to the BOLD effect highlights EEG’s sensitivity to rapid, preconscious dynamics that precede conscious report.

Direct EEG–fMRI coupling further supports this account. Across undisturbed rest (first 500 TRs/session), the instantaneous amplitude of the *α* spatio-spectral pattern was inversely associated with voxel-wise BOLD in bilateral parietal/occipital cortices (Fig. 5A), replicating prior work [13]. Spatial correlation with 14 RSNs confirmed the strongest overlap with the high-visual network (Fig. 5B), reinforcing that low *α* co-occurs with enhanced sensory processing and vice versa.

EEG-based classification was moderate. This likely reflects (i) conceptual heterogeneity in broad internal/external labels that a single spectral feature cannot fully resolve, and (ii) measurement constraints of EEG in MRI environments (residual pulse and gradient artefacts) despite advanced preprocessing [13, 77–79]. Future EEG-only work outside the scanner could improve signal fidelity and ecological validity, and additional markers (e.g., connectivity, cross-frequency coupling) may increase separability [77]. Nonetheless, the present EEG results provide temporally precise evidence that *α* oscillations track spontaneous shifts in attentional orientation.

### 3.3 Experience sampling and the phenomenology of attention

Descriptive Experience Sampling (DES) moves the categorization task from participants to trained researchers. This allows high standards of ecological validity in obtaining data about experiential states and offers a powerful bridge between first-person and third-person perspectives [51, 80], but at the same time, DES introduces interpretive challenges for dichotomous distinctions such as internal vs. external awareness orientation.

First, DES’s richness requires theory-driven reduction. As in our precursor study [24], many samples were *hybrids* (e.g., concurrent perception and imagery) that resist binary assignment. Such hybridity suggests that attentional orientation spans a continuum rather than a switch [54, 81]. While we used forced coding to enable analysis, the presence of hybrids motivates refined taxonomies and analysis tools that capture graded, overlapping modes of consciousness (see Supplementary Material 1).

Second, DES’s focus on ecological validity of undisturbed moments complicates cross-study comparison. Our paradigm (and 24) uses long (5 min) intervals between samples; many ES protocols probe frequently, approximating a monitoring task [24]. This methodological difference may explain the absence of DMN differences here: participants may have been mind-wandering even during “external” moments. Yet, as a previous DES-fMRI study [24] explicitly found DMN, as a hallmark of internally oriented states, it is necessary to point out possible variability in DES training and facilitation of interviews [82], participant heterogeneity, and variability in coding criteria may also contribute. Automated or AI-assisted coding could improve reproducibility and support open repositories of mind–brain correspondences.

A temporal nuance is also notable. Although DES anchors reports to “the moment before the beep,” our fMRI effects extend over longer pre-beep intervals, whereas EEG effects are confined to 1–2 s. This suggests a hierarchical temporal architecture, whose parts are captured separately across modalities. State orientation (internal vs. external) may persist over longer neural timescales, while content fluctuates rapidly. A layering model could reconcile differences between modalities and sampling schemes, and is consistent with recent models proposing that intrinsic neural timescales vary across cortical hierarchies—shorter in sensory regions and longer in associative and transmodal areas [83, 84]. Simultaneous EEG-fMRI studies have begun to disentangle these nested timescales, revealing distinct but complementary contributions of fast electrophysiological signals and slower hemodynamic fluctuations to spontaneous cognition [85–87].

### 3.4 Limitations and Future Directions

While this study offers novel insights into the interplay between spontaneous experience and largescale brain dynamics, several limitations should be acknowledged.

First, the use of Descriptive Experience Sampling (DES) inherently relies on subjective introspection, which, despite rigorous training protocols, may still introduce variability across participants. Although this subjectivity is a strength in capturing phenomenologically grounded data, it also presents challenges for replicability and generalizability [88, 89].

Second, the integration of EEG and fMRI data, while powerful, is constrained by the differing temporal and spatial resolutions of these modalities. The fusion of fast electrophysiological signals with slower hemodynamic responses complicates direct inference, especially in dynamic contexts such as spontaneous thought [90, 91]. Future work may benefit from improved computational models or hybrid acquisition protocols that better align these signal domains [92].

Third, the sample size was limited, reflecting the intensive nature of multimodal DES protocols. Larger, more diverse samples will be needed to confirm the generalizability of these findings and to assess individual differences in spontaneous mentation more robustly [93].

Looking forward, combining DES with realtime neural decoding methods or ecological momentary assessment could help move beyond post-hoc correlations toward closed-loop, interactive models of conscious experience. Additionally, applying this framework in clinical populations—such as those with depression, ADHD, or schizophrenia—may yield mechanistic insights into altered self-generated thought and its neural substrates [94, 95].

## 4 Conclusion

Taken together, our results situate spontaneous thought and attention as fundamental components of adaptive cognition. The brain’s intrinsic dynamics appear to oscillate between *external readiness* (multisensory recruitment, salience–attention coordination) and *internal reflection* (suppression of sensory inflow, self-generated mentation) [93, 94, 96]. This rhythmic interplay has been proposed to reflect a functional balance between the default mode network (DMN) and task-positive networks such as the dorsal attention and salience systems [97, 98]. Evidence from dynamic functional connectivity studies suggests that the brain transitions through temporally evolving configurations that preferentially support externally oriented processing or internally generated thought [90, 91], while phenomenological evidence from descriptive experience sampling supports the notion that these modes can also cooccur within a single moment of experience[24]. Conceptually, this aligns with views that resting cognition reflects an active readiness to engage with the environment while also supporting the internal updating of generative models when sensory input is attenuated [99, 100].

The methodological contribution is equally important. Integrating DES with simultaneous EEG–fMRI and machine learning provides a template for uniting subjective experience with objective brain dynamics in naturalistic contexts [89]. Beyond basic science, this approach has implications for domains in which balance between internal and external focus is critical: creativity, learning, and clinical conditions marked by altered self-generated thought [90]. By directly linking phenomenologically grounded experience to multimodal neural signatures, we move beyond inference from task contrasts toward time-resolved accounts of how large-scale networks and oscillations shape the ebb and flow of human experience.

Ultimately, our findings frame spontaneous mentation not as a by-product of an idle brain but as an organized and measurable expression of the mind’s natural dynamics. Bridging neural and experiential data brings us closer to an empirical science of consciousness that honours both biological grounding and subjective richness—an ambition traceable to William James and increasingly realizable with contemporary multimodal neuroscience [96] employing integrated measurement designs[101].

## 5 Methods

### 5.1 Participants and study design

We collected a data set composed of a series of simultaneously recorded physiological (EEG, fMRI), behavioral (eye-tracker), and phenomenological measures while the participants were in a resting-state condition. The complete data set comprises simultaneous EEG-fMRI time series as well as several structural MR images. The data were acquired during a 7-year period at the National Institute of Mental Health (NIMH) in Klecany, Czech Republic. For this specific study, we excluded all participants’ eye-tracker recordings because this part of the data set wasn’t complete for all participants. The study comprised eight healthy volunteers (mean age: 30.1, range: 20 - 45, 2 males and 6 females). Within the study, they are identified by pseudonyms. With the exception of the additional EEG and eyetracker modalities, the data acquisition process closely emulated the methodologies employed in Hurlburt et al. [43]. Prior to image data acquisition, each participant underwent four to six training sessions with four to six sampling events in their natural everyday environment to train themselves to apprehend momentary experiences in accordance with the Descriptive Experience Sampling method [51] and get accustomed to its procedure. Each training collection session was followed within twelve hours by an expositional interview session with a researcher, where the samples were processed and where the proper training happened. These training samples were not used in the study. The core of the training was the ability to apprehend a pristine experience happening in the last uninterrupted moments before the beep and to make hand-written notes right after apprehension as a basis for a sample-collecting interview. Within weeks after the training, the participant underwent the 9 sessions of 25-minute eyes-open EEG-MRI recordings with an eye-tracker. The instructions for the MRI session were similar to typical restingstate recording: “Please relax without falling asleep, and do keep your eyes open”. These sessions contained 4 pseudo-random beep events with a mean interval of 5 minutes, where participants apprehended their “before the beep” experiences and made notes about them without the use of sight. These notes were immediately processed after each session in an expositional interview. A participant usually completed two such sessions in a day, resulting in 5 to 6 visits. After the final participant session, we created a dataset of textual descriptions of experience samples from the recorded interviews. The study was approved by the ethical committee of NIMH and was conducted in accordance with the Declaration of Helsinki. All participants provided written informed consent to participate in the study.

### 5.2 Data acquisition

The imaging was performed using a 3T Siemens Prisma magnetic resonance scanner with a 64-channel RF receiving head coil (Siemens Healthineers, Erlangen, Germany). The functional imaging (fMRI) was performed using a multiband T2*-weighted multi-echo 2D echoplanar imaging (EPI) sequence with the following parameter: multiband factor (MB) = 4, repetition time (TR) = 700 ms, echo time (TE) = 30 ms, 48 axial slices, acquisition matrix = 74 *×*74, field of view (FOV) = 222 *×*222 mm, isotropic resolution = 3*×* 3 *×*3 mm^3^, flip angle (FA) = 52 °. The structural MR images were acquired using a T1-weighted magnetization prepared rapid gradient echo (MPRAGE) sequence with parameters: repetition time (TR) = 2300 ms, echo time (TE) = 2.33 ms, 240 sagittal slices, acquisition matrix = 224*×*224, field of view (FOV) = 224*×*224 mm, isotropic resolution = 1× 1 ×1 mm^3^, flip angle (FA) = 8 °.

The EEG data with a sampling frequency of 1000 Hz were recorded with an EGI Hydrocell MR-compatible 256-channel high-density electrode net plugged into the EGI GES 400 signal amplifier (Electrical Geodesics, Inc., Eugene, Oregon, United States). Two extra electrodes were used to acquire electrocardiogram (ECG) during the recording in the MR scanner, providing a reference signal for the removal of the ballistocardiogram artifact. EEG acquisition was accurately synchronized with the MR scanner using the GES Clock Sync I/O device, ensuring precise temporal alignment between the EEG system and the MR scanner. Additionally, TR triggers for gradient artifact removal were recorded via the GES Clock Sync I/O.

The EEG net was applied in a preparatory room, with electrode impedances adjusted to ensure that all electrodes exhibited an impedance below 60 kΩ. Both EEG and ECG signals were checked for data quality. Afterward, the subject was disconnected from the EEG amplifier, positioned in the MR scanner, and reconnected to the EEG amplifier. Data transfer was facilitated through an optical fiber connection.

The entire screen was set to white throughout the session (except for the initial instruction screen) to provide natural light conditions for note-taking.

A ten-second long square waveform at a frequency of 1kHz was used as a signaling sound. Auditory stimuli were presented using MR Confon HP AT 01 insert earphones with 300 mm air tubes and foam ear pieces that utilized the permanent magnetic field B0 as a magnet for electro-mechanical transduction. The transducers were positioned near the head coil, and the sound was delivered to the ears through tubes functioning as waveguides. A low-frequency amplifier (MR Confon - Starter f MKII+ amplifier) was used to adjust the auditory stimuli to an appropriate sound level. The sound level of the auditory stimuli was determined experimentally in collaboration with the test subject prior to the first DES session. A comfortable level ensuring clear signal recognition was identified and maintained consistently across all subsequent sessions.

Both visual and auditory stimuli were generated using a custom presentation script implemented in E-Prime 2.0 software. The beginning of the presentation script was synchronized with the first TR trigger delivered through the GES Clock Sync I/O. The visual and auditory events were synchronized with EEG acquisition using the Experimental Control Interface (ECI) protocol. As a result, both TR triggers and experimental event triggers were included in the final EEG recording, enabling integration of all modalities.

### 5.3 MRI data preprocessing

Prior to the analyses, the functional images were preprocessed using a standard preprocessing pipeline in the CONN Toolbox (https://web.conn-toolbox.org/) for Matlab (The MathWorks, Inc., Massachusetts, USA) labeled as ‘default mni’. The preprocessing steps consisted of realignment and unwarp (motion correction); slice-timing correction; outlier identification; direct segmentation and normalization to the MNI space; and spatial smoothing with an 8 mm full-width half maximum (FWHM) kernel.

### 5.4 EEG data preprocessing

To derive the *alpha* spatio-spectral pattern, our approach closely aligns with the methodology employed in our previous study [13]. Therefore, raw EEG data (whole recordings) were preprocessed by a fully automated pipeline introduced in [102]. The preprocessing pipeline utilizes builtin and in-house MATLAB (MathWorks, Natick, MA, United States) functions as well as SPM [103], Fieldtrip [104], and EEGLAB [105] toolboxes. The preprocessing steps are summarised in the following paragraph.

The first step was gradient artifact removal by the FMRI Artifact Slice Template Removal (FASTR) method [106] in EEGLAB followed by a ballistocardiogram artifact removal utilizing adaptive optimal basis set method introduced in [78]. Identification of bad channels relied on criteria such as low correlation across all channels in the wide frequency band (1 - 80 Hz) and variance in the EEG non-physiological frequency band (200 - 250 Hz). If any criterion marked a channel as an outlier in the distribution across channels, the channel time course was interpolated using the time courses of neighboring channels. Subsequent steps involved filtering EEG data within the 1 - 80Hz frequency band and applying Independent Component Analysis (ICA) to eliminate movement and other biological artifacts, including electrooculographic (EOG) and electromyographic (EMG) artifacts. FastICA [107] algorithm, employing a deflation approach and hyperbolic tangent as the contrast function, was utilized. Artifactual components were identified based on three parameters: correlation values between ICs and reference EOG and EMG signals, the similarity of ICs power spectrum with a 1/f function, and kurtosis of ICs timecourse [108]. Despite comprehensive preprocessing, we opted to exclude cheek electrodes and the two lowest layers of neck electrodes from further analysis, in line with concerns in current literature about disproportionately higher artifact contamination at these electrode sites [109]. Consequently, 195 out of 257 electrodes were retained for subsequent analyses. As a final preprocessing step, the EEG data were re-referenced to an average reference.

### 5.5 Descriptive Experience Sampling and coding of samples to two target groups

Descriptive experience sampling is a method of facilitated introspection. It is an idiographic procedure aiming to acquire directly apprehended (“pristine”) inner experience [51] in the form of high-fidelity textual natural language descriptions. Its essence characterizing it among other phenomenological approaches is by its authors summed up to four overlapping methodological characteristics [44]: 1) limiting the investigation to specific and clearly identified moments; 2) limiting the experience to its pristine qualities (limiting to direct apprehension); 3) bracketing presuppositions and 4) iterative acquirement of skill. It institutes data acquisition conditions with both constraining and supporting scaffolding, which can form a data processing “pipeline” of its kind. The procedure starts with the participants’ apprehension of the experience sample on a familiar sound signal, where their focus is guided by an open and in-training familiarised question: “What was in your experience at the moment of the beep?”. The start of the signal marks the immediate moment before it as a target for the sample, while the length of the beep (10s) serves as a “freezing” state for capturing the experience in working memory. Then participants write quick notes about the experience and return to the contextually natural activity. In the resting-state EEG-fMRI 25-minute-long session, participants undergo 4 such events. After each such session, the written notes are the basis for an expositional interview, where the participant and researcher have an open conversation using the four DES principles about each sample. The apprehended content of the sample supplemented by participants’ recollection is processed during the interview towards the orally expressed high-fidelity descriptions, and later from the recordings summed up to written form.

Three researchers familiar with the previous DES study about internally and externally oriented experiences [24] then read the dataset of samples separately and forced-categorized them into two groups. This was accompanied by the mark of confidence in the validity of the categorisation (interval 1-10, 10 being the most valid). The confidence scale was not further utilized within this study. The final dataset of samples with externally and internally oriented experience was constructed from the full agreements in the same category.

We have collected 288 DES samples in total (8 participants x 36 samples), while in the data analysis, we entered 240 of them (42 samples did not achieve coder agreement, and 6 samples were removed because their corresponding brain data were marked as outliers). Of these 240, 163 were categorised as predominantly internally and 77 as predominantly externally oriented experiences.

### 5.6 BOLD features

To capture the brain response by fMRI, we used 14 large-scale resting-state brain network masks by Shirer et al., 2012 [53], see Fig 1A as regions of interest (ROIs). We opted for these masks as they cover the main brain networks that are generally recognized by the community. From each individual fMRI session, we extracted the corresponding time series averaged across the voxels in each ROI. The extracted time series were linearly interpolated to match the sampling frequency of the EEG signal. The resulting time series were normalized to z-scores.

To identify the samples that exhibited an a typical response to beep, we calculated an average response of the 14 brain networks to the beep across all 288 samples in a window that stretched from the moment of the beep to 25 seconds after the beep. A typical beep response was represented by the first principal component across the 14 brain networks. We then calculated Spearman’s correlation coefficient between the individual beep responses from the corresponding time window and the typical beep response. In total, 39 BOLD samples with *r <* (*Q*1 −0.5 *×* (*Q*3 −*Q*1)) were marked as outliers.

To synchronize the time-locked phenomenological samples with the response in the BOLD signal, we had to compensate for the delay, which is an inherent property of the BOLD signal. The BOLD signal reflects the regional hemodynamic changes and the ratio of the oxygenated and deoxygenated blood in brain vessels, and as such, it represents a proxy measure of the actual neuronal activity [110]. Due to the hemodynamic response, the peak in the BOLD signal that we observe is delayed for ca. 4-6 seconds after the actual neuronal activity [111]. As we focused on the subject’s inner experience preceding the moment of the beep, we attempted to estimate the actual delay of the hemodynamic response from our data and adjusted the times of the beeps accordingly. We estimated the delay of the hemodynamic response by finding the beginning of the leading edge of each subject’s average BOLD signal from the 14 brain networks by Shirer et al. [53]. The leading of the signal was found as the first statistically significant activation (using Wilcoxon rank sum test *p <* 0.01) above the baseline. The baseline was created from time-averaged 4.5 minute long time courses of the 14 brain networks before the moment of the beep. We then used the estimated delay, averaged across subjects, and adjusted the times of the beeps by -3.1 seconds.

### 5.7 EEG features

To derive the *alpha* spatio-spectral pattern, we generally followed the implementation of the electrode space version of the spatio-spectral decomposition method from [13]. The approach assumes that the electrical activity expressed as the EEG signal envelope measured at the electrode locations on the scalp is a linear mixture of source signals representing the brain activity. We further assume that the brain electrical activity may have a different spatial profile across different EEG frequency bands. At first, we extracted 60 seconds of EEG recordings preceding each valid beep. We applied a short-time Fourier transform (STFT, 1000 ms window size, 100 ms window step) on each time series and extracted BLP time series for pre-defined frequency bands: Delta (*δ*, 1 - 4 Hz), Theta (*θ*, 4 - 8 Hz), Alpha (*α*, 8 - 12 Hz), low Beta (*β*_1_, 12 - 16 Hz), middle Beta (*β*_2_, 16 - 20 Hz), high Beta (*β*_3_, 20 - 30 Hz), and Gamma (*γ*, 30 - 40 Hz), resulting in 7 BLP matrices (one for each frequency band). To ensure that the resulting BLP time series are not influenced by the brain activity after the beep, we cut the end of each time series by 500 ms. Next, we concatenated BLP matrices of all frequency bands along the spatial domain, thus forming a joint spatio-frequency domain on the single-session level. Each individual session BLP time series was then z-score normalized to handle inter-session variability. All single-session normalized BLP matrices were subsequently concatenated along the temporal domain forming a group BLP matrix. Applying a temporal (group-level) ICA by RUNICA algorithm [112] with a PCA dimensionality reduction set to *C* = 30 components (based on a typical number of components recovered in BOLD resting-state ICA analyses), we obtain 30 independent components (ICs) representing spatio-spectral patterns. Since ICA decomposition is based on iterative algorithms with random initialization, we utilized the ICASSO tool [113] to investigate the algorithmic stability of the ICA decomposition itself by performing the ICA decomposition 20 times with different initial conditions. The cluster’s centroid time series were then considered as the most reliable estimate of components and, therefore, utilized for the subsequent analyses. For more methodology details, please refer to [13]. We visually identified two spatio-spectral patterns resembling the *alpha* pattern from [13]. We utilized the ICA mixing and PCA weights matrices to obtain weights in the original spatio-frequency domain for obtaining the final spatio-spectral signature (based on corresponding rows and columns of the two selected components). The final EEG feature time series was obtained by correlating each time sample of the full BLP matrix with the spatiospectral signature. Therefore, the EEG feature can be interpreted as a similarity of the current spatio-spectral BLP profile to the spatio-spectral signature. The feature time series defined this way was further utilized in classification tasks and exploration in terms of pattern I/E time series.

### 5.8 Classification task

As each DES resting-state session contains four quasi-randomly placed beeps, with nine sessions per subject, we obtained 288 pre-beep samples in total. In total, 48 samples were excluded from further analyses due to reasons stated in the section above. This resulted in 240 samples and 14 predictors.

Although each modality data was used in a separate classifier, the following feature preparation procedure applies to both. We assumed that the experience should be distinguishable somewhere in the last 5 seconds prior to the beep, therefore, we extracted the data points from each modality from -5 seconds before the beep up to -0.5 seconds before the beep, which (with a 0.1-second step) resulted in 46 samples. As the exact position of the subject-individual neural correlates in time was unknown and naturally slightly different between subjects, we decided to incorporate temporal averaging of each modality signal. Using a window size that ranged from 0.1 seconds (with a 0.1-second step) up to 2 seconds, we created 20 variants of temporally averaged original signals. Eventually, we extracted features from 730 combinations of window positions and window sizes 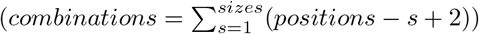.

To classify the extracted features, we used a support vector machine (SVM) classifier implemented in Matlab (The MathWorks, Inc., Massachusetts, USA). Due to class imbalance (163 internal samples vs. 77 external samples), we adjusted the misclassification cost weights that penalize the misclassified samples. The misclassification cost for the larger class (I) was set as a ratio of the number of E and I samples (77/163). To evaluate the classification model and prevent it from overfitting, we used a leave-one-out cross-validation technique. The classification of I/E samples using a linear SVM was performed for all combinations of 20 window sizes and 46 window positions, resulting in a 20 *×* 46 matrix containing the classification accuracy values of the 240 samples in total, see Fig 2A,C. To test the achieved classification accuracy, we compared our results against a null distribution generated by performing 5000 permutations of the I/E labels (separately within each subject) for each of the 20 × 46 combinations of window sizes and positions. Eventually, we used a threshold-free cluster enhancement (TFCE) approach [114] with a height parameter *H* = 2.0 and an extent parameter *E* = 0.5 to evaluate the clusters in the original matrix as well as in the permuted data.

### 5.9 EEG alpha spatio-spectral pattern relation to BOLD

We computed an fMRI BOLD correlate of *alpha* EEG spatio-spectral pattern to directly link both modalities in this study. To not include any beep event (sample of experience), we extracted the first 500 TRs BLP time series for each session. We utilized the EEG feature spatio-spectral signature as a filter for the first 500 TRs session spatio-spectral BLP time series. Specifically, we computed a dot product between each time sample of single-session spatio-spectral BLP matrix and EEG feature spatio-spectral signature. Those single-session time series were further downsampled to match the BOLD TR and band-pass filtered (0.008 - 0.09 Hz), which is a typical step for resting-state fMRI analysis. To assess the single-session relationship between EEG spatio-spectral component time series and the BOLD signal in each voxel, the general linear model (GLM) was utilized.

As in [13], we included three regressors into the general linear model for each component separately: a component time series convolved with the canonical hemodynamic response function as a first regressor, a component time series convolved with the first (temporal), and the second (dispersion) derivative of the canonical hemodynamic response function as proposed and examined in event-related studies [115]. To capture the sign (positive/negative correlation) of the relationship between component BLP time series and the BOLD signal, we implemented a formula proposed in [116] to aggregate all three regressors to a single beta coefficient which was further utilized. Group-level statistics was performed by a one-sample t-test at each voxel. Cluster-based permutation statistics [117] was applied to correct for multiple comparisons (*α* = 0.01, *α*_*cluster*_ = 0.05). The statistical GLM map was visualized with the BrainNet Viewer tool [118].

Moreover, we examined the correlation between the spatial statistical map mask derived from Figure 5A and each of the 14 functional networks employed in extracting BOLD features (refer to Figure 5B). This aspect of the analysis was not subjected to statistical significance testing.

## Acknowledgments

This project was supported by the ERDF-Project Brain dynamics, No. CZ.02.01.01/00/22 008/0004643 (DT, SJ, VK, LJ, JH), the Czech Technical University Internal Grant Agency - grant number SGS23/120/OHK3/2T/13 (SJ) and the Czech Science Foundation project No. 21-32608S (TH, DT, SJ, VK, LJ, JH). This publication was written with the support of the Johannes Amos Comenius Programme (P JAC) provided by MSMT reg. n. CZ.02.01.01/00/23 025/0008715 (TH, JH, LJ).

## Data and Code Availability Statement

The datasets (preprocessed features of fMRI and EEG modalities) and classification pipeline used for analysis during the current study will be made publicly available on a public GitHub repository upon publication of this manuscript.

## Supplementary information

Supplementary File provides five detailed observations illustrating the hybridity of internality and externality in participants’ experience samples [24].

## Notes

### Competing Interest Statement

The authors have declared no competing interest.

