## Supplementary Material 1 for "Spontaneous thought orientation tracked by fMRI networks and EEG alpha power dynamics"

### Five observations about the hybridity of internality and externality in the experience samples

We have corroborated all five of Fernyhough et al. (2018) observations, which suggest the existence of experiential hybrids of combined internal and external attention and which express concerns about the strict switching model, holding that the existence of an internal train means full external train suppression:

1) Our participants reported experiences with simultaneously external visual attention to the environment and internal images: e.g., "Jane saw a part of the face of a person in the MRI control room in the mirror, at the same time she saw a scene from childhood memory - she was lying on the bed and saw other children around her. The internal image was the focus of her experience, but external sensation was also present."

2) They purposefully manipulated external experience trying to suppress it or enhance it, e.g. "Anna sees part of the fMRI tunnel, in which she imagines a simple human-toy-like figure with the head and upper body in the shape of letter T; she can see partially face of the figure and its emotional state ("mischief") from its gray eyes looking to the right; this image is understood as a product of her purposeful imagination, but it is not effortfully imagined in the moment of the beep." Within describing the context of apprehension, Anna described that the image of the T figure was produced as a part of playful manipulation within her visual field.

3) We have captured in their experience moments of internally guided action (intent of the action) connected with external sensation: e.g., "George relaxed fingers on his feet and felt relief in them, at the same time he felt worried from actively moving too much; the worry was connected with constraining feeling of responsibility for not moving in the MRI. He described the feeling as active overseeing of his motor control."

4) There were cases of participant's attention to the external environment, which contains external sensation with layers of an internal supplement of added meaning associated with the sensations: e.g. "Anna is actively seeing border of the screen, and pondering what is behind the screen, at the same time she experiences agitation and panic from the feeling, that what sees is not real. She classifies the feeling as derealisation."

5) In the coding of the experiences of bodily sensations, we also had to create an arbitrary border between internal and external experiences in making bodily sensations external: e.g., "Case perceived a sensation of cold in his feet." was coded as an externally oriented experience.

(Names of participants are anonymized; identification uses the same pseudonyms as the main text.)
